# Scale Invariance of Mechanical Properties in the Developing Mammalian Retina

**DOI:** 10.1101/2024.10.21.619491

**Authors:** Elijah Robinson Shelton, Michael Frischmann, Achim Theo Brinkop, Rebecca Marie James, Lucas Maria Hüttl, Friedhelm Serwane

**Affiliations:** Faculty of Physics and Center for NanoScience (CeNS), Ludwig Maximilian University, Munich, Germany; Institute for Biophysics, University of Ulm, Ulm, Germany

## Abstract

The computational capabilities of the human central nervous system arise from neuronal architectures. Neuronal tissues are built through force-driven physical remodeling over durations ranging from seconds to days. However, how cell-generated forces accumulate and relax to drive neurogenesis remains unknown due to the difficulty of applying stresses (i) in developing mammalian nervous tissue, (ii) directly within the cellular microenvironment, and (iii) for durations spanning seconds to hours. Previous studies have shown that non-neuronal tissues and cells in 2D culture remodel in scale-free manners. Whether and how this translates to developing neuronal tissues remains an open question.

Here, we probed the mechanics of mammalian neuronal tissue on developmental timescales. We accessed developing mammalian neuronal tissue using retinal organoids, stem-cell-derived models of the retina. To probe mechanics at scales relevant for retinal development, we used magnetic droplets as long-term mechanical actuators. We recorded strain responses to applied stresses across four orders of magnitude in time, up to one hour. We found that dynamic creep compliance and tensile moduli follow a power law with an exponent consistent with a material just above the glass transition. This scale-free rheology represents an unprecedented description of nervous tissue mechanics on developmental timescales and opens the door to a biophysical understanding of the emergence of functional neuronal architectures.

## I. INTRODUCTION

How the architecture of neuronal tissue establishes its electrical functionality is one of the big open questions of neuroscience[1]. To develop the central nervous system (CNS), mechanical forces work together with biochemical factors to drive the structural remodeling necessary to build a synaptically connected network[2–4]. The retina is a prime example, transforming from a single-layered neuroepithelium into a multilayered, light-sensitive neuronal tissue. It has become a focal point for researchers investigating the links between molecular factors, cellular forces and tissue organization[5, 6]. The self-organized wiring up of the layered neuronal architecture during retinogenesis is inextricably linked to tissue mechanical properties. Mechanical cues from the microenvironment drive differentiation, and its rheological properties determine the physical demands of neuronal translocation required for cells to arrive at their designated position within the tissue. However, in humans and other mammals, it is experimentally difficult to quantify the rheology underlying neurogenesis as this requires (i) accessing mammalian nervous tissue, (ii) applying a known force directly in the microenvironment in which cells undergo neurogenesis, and (iii) sustaining the controlled mechanical interrogation over the long timescales relevant to neuronal translocation and integration of mechanical signals, for seconds to hours.

A deep interest in the physics of the nervous system has inspired researchers to characterize its mechanical properties in the regime of shorter timescales using conventional tools. Elastic responses have been measured on the second timescales and reported for living guinea pig and ruminant retina slices using the Atomic Force Microscope (AFM)[7, 8], and the mechanics at nanosecond actuation periods have been mapped for the mouse retina using Brillouin microscopy[9]. Discrepancies between AFM and Brillouin microscopy measurements have been observed and could be attributed to differences in length and timescale of the mechanical stimulation, highlighting the key role of timescales in mechanobiology research[10]. At the subcellular scale, researchers have quantified the time dependencies of mechanical properties using a plethora of techniques[11, 12]. For instance, probing the short term mechanics using optical tweezers repeatedly over several days could reveal the role of elastic moduli for developing spheroids[13]. However, conventional methods like the AFM or optical tweezers have not met the experimental challenge of continuously applying a controlled force over an hour while measuring the strain response in the presence of morphogenetic movements. In the case of the retina, and neuronal tissue in general, measurements of mechanical properties on the minutes to hours timescales, which are critically needed to understand the physical context of neurogenesis, are currently lacking[4, 14].

Recently, stem cell-derived *in vitro* organoid models have provided researchers with systems which complement traditional animal models as platforms for studying the details of retina development. Termed retina organoids[6, 15], they display all cell types and mimic the tissue architecture of the *in vivo* retina[16–18]. In both *in vitro* and *in vivo* systems, retina development starts with an ensemble of neural progenitor cells derived from embryonic stem cells (ESCs)[16, 19]. Parts of the ensemble transform into a single layer neuroepithelium which then becomes the neural retina[16, 19]. At this stage, researchers have measured the elastic properties of its surface using an AFM on mouse retina organoids[16]. The results have served as inputs for mechanical modelling using vertex models[16, 20, 21] and successfully explained the morphological changes the neuroepithelium undergoes before it starts converting into a 3D laminated tissue. Observations within this pseudostratified epithelium both *in vivo* and *in vitro* have found that neural progenitors undergo interkinetic nuclear migration (IKNM) between cell divisions[6]. After a period of proliferation, cells exit the cell cycle, differentiate, migrate into the correct radial and lateral positions, and form synapses[22–24].

In contrast to previous experiments, we reveal the mechanical properties which contextualize the neuronal movements at scales of neurodevelopment by probing inside living mammalian retina tissue. By combining retina organoids as *in vitro* models for the mammalian retina and magnetic droplets as *in situ* mechanical actuators[25], we were able to measure the time-dependent mechanical properties on timescales from seconds to hours, and carry out mechanical characterizations over multiple days of development. Revealing the long-term mechanics associated with neural development required three key advancements in the use of magnetic droplets for measuring tissue rheology: First, we applied stresses for up to one hour, extending the duration by a factor of four compared to previously reported measurements[26–28]. Second, we calibrated stresses directly in viscous standards, a simplified approach which we validated against a shear plate rheometer with both viscous and viscoelastic samples, and made these tools available to the community (see Methods). Third, we employed a conceptually different biophysical model which does not contain any intrinsic scales in contrast to previous reports[26–28]. Previous models performed well within a limited observation window but diverged from the data near the edges of the measurement range (Suppl. Fig. 4). In contrast, our model maintains high fidelity across the entire measurement window, indicating that the extracted mechanical properties accurately reflect the mechanics of the system throughout the observation period. This approach allowed us to reliably extract mechanical properties not only during the central portion of the measurement window but also near its boundaries. With these distinct advancements, we were able to capture the biophysics of neurodevelopment across several orders of magnitude in time.

## II. RESULTS

### A. Probing long-term mechanics in a mammalian retina organoid

To address the challenge of accessing developing neuronal tissue, we cultured an *in vitro* model of the mammalian retina following an established protocol for generating retina organoids from mouse embryonic stem cells (ESCs) (Fig. 1a; Methods)[16, 29]. The Rx-GFP line provides a fluorescence-based readout for the differentiation into retina cell types as GFP is expressed under the retina tissue specific promoter Rx[16]. We confirmed development of photoreceptor cells by recoverin immunostaining and confocal microscopy imaging in organoids from the first round of culture in which we collected mechanical measurements (Fig. 1b; Suppl. Movie 1, Methods).

**FIG. 1:**
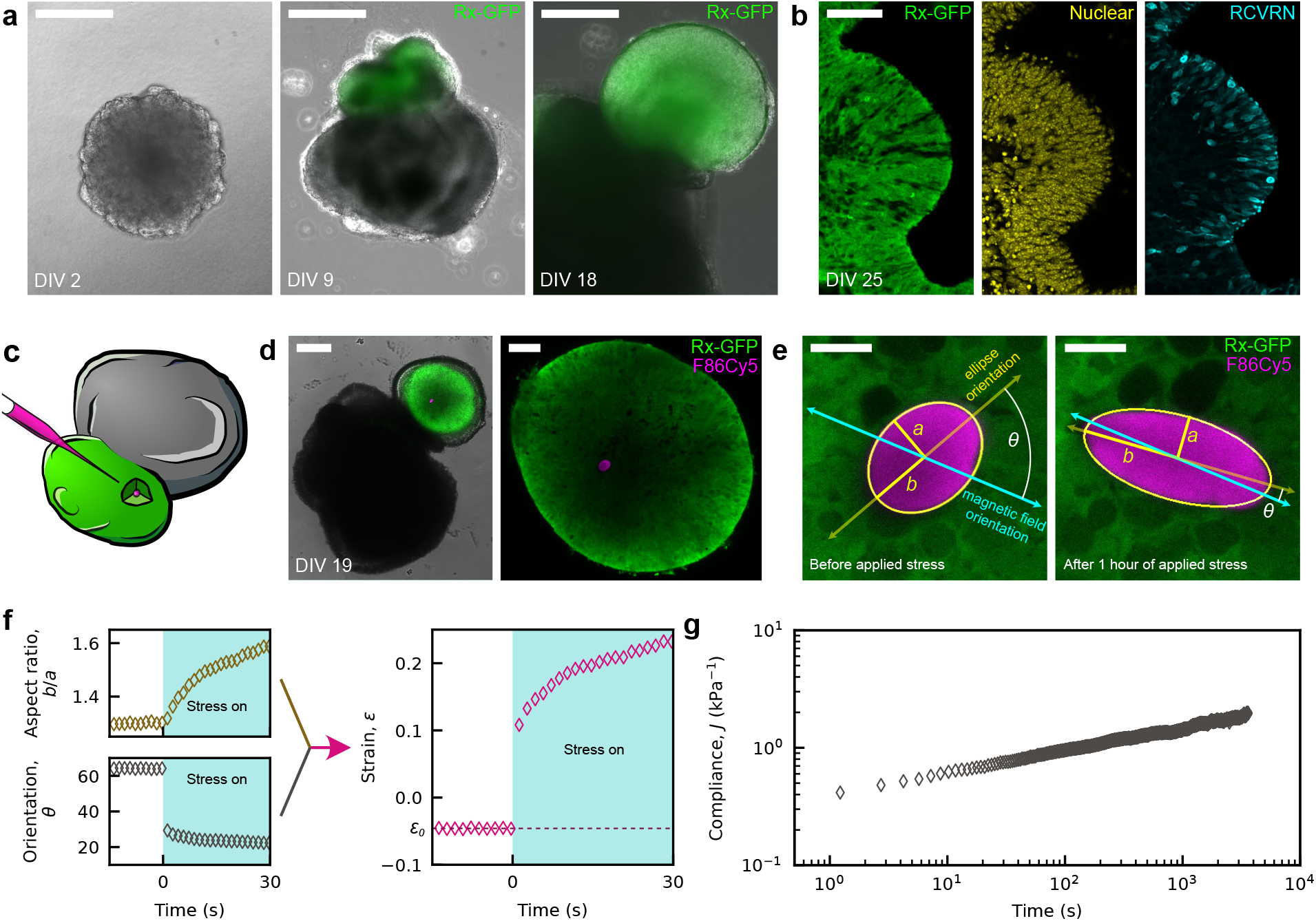
Probing the mechanics of mammalian retina organoids *in situ* using ferrofluid droplets as mechanical actuators. **a**, Phase contrast and fluorescence microscopy images of retina organoids grown from mouse embryonic stem cells at days in vitro (DIV) 2, 9, and 18. Retina cells become visible as GFP is expressed under retina promoter Rx at DIV 9 and 18. **b**, Confocal microscopy images of a cryosectioned organoid fixed on DIV 25 showing, left to right, GFP, nuclear stain, and recoverin immunostain (See Suppl. Movie 1). **c**, Sketch showing the injection of fluorescent magnetically responsive oil droplets into retina tissue. **d**, Confocal and transmitted light images of a DIV 19 retina organoid with magnetic droplet after injection. **e**, Confocal microscopy images of droplet (magenta) embedded in retina tissue (green) before (left) and after (right) the onset of applied stress (See Suppl. Movie 2). The elliptical segmentation of the droplet is shown (yellow), with the lengths *a* and *b* of the minor and major semi-axes and the relative orientation of the ellipse, *θ*, with respect to the magnetic field. **f**, Plots of *b/a* and *θ* versus time demonstrate how the aspect ratio and orientation change with onset of stress (left). Together, *b/a* and *θ* determine the strain *ε* (right). Dashed line indicates initial strain *ε*_0_. **g**, The compliance *J* = (*ε − ε*_0_)*/σ* is plotted against time for the entire duration of applied stress (1 hour). *σ* = (371 *±* 12) Pa (mean *±* s.d., see Methods). Scale bars: **a** 200, **b** 50, **d** 200 (left), 50 (right), **e** 10, all in microns.

To meet the demand of interrogating inside the the neuronal tissue, we used ferrofluid microdroplets to probe the rheology of the microenvironment experienced by neural progenitors. We applied a carefully calibrated stress dipole to the retina tissue using magnetically responsive droplets, building on our previous experiments in non-neuronal systems[25, 26], as detailed in Methods. Previous implementations of the technology relied on the measurement of the ferrofluid magnetization curve as well as the magnetic field. Both required specialized equipment such as a superconducting quantum interference device (SQUID)[25]. Here, we calibrated the applied stresses using only Newtonian fluids with known viscosity as a standard, making the technology accessible to other researchers (Methods). To improve analysis, we developed a python-based open source software package for data analysis with a graphical user interface (GUI) and made it available to the community[30], (Methods, Suppl. Fig. 1). To reliably quantify tissue strain along arbitrary directions within the imaging plane we optimized the strain readout and image segmentation (Methods, Suppl. Note 1). Taken together, these improvements allowed us to perform hour-long compliance measurements with high fidelity. We injected droplets with radii between 9 and 26 microns which were both magnetic and fluorescent[31] into retina regions which we identified via the fluorescence signal (Fig. 1c and Methods). Following an injection, we incubated the organoid for several hours and then performed live-imaging (Methods). For each measurement, we confirmed that the droplet was situated inside the thin (up to 100 microns) retina layer (Suppl. Movie 1) using confocal microscopy (Fig. 1d). We collected time series of confocal sections before and during the application of stress (Fig. 1e, Suppl. Movie 2, Methods), and quantified the droplet’s aspect ratio *b/a* and relative orientation *θ* at every time point using an elliptical segmentation (Fig. 1e,f, Methods). If the major axis of the droplet is aligned with the magnetic field (*θ* = 0), then the strain is a function of only the aspect ratio: 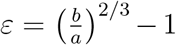. In general, the droplet and magnetic field may be misaligned, as other forces in addition to those from the magnets are present in the tissue. Therefore, we calculated the strain along the direction of the magnetic field from the aspect ratio and orientation of the droplet in the following way[32]:

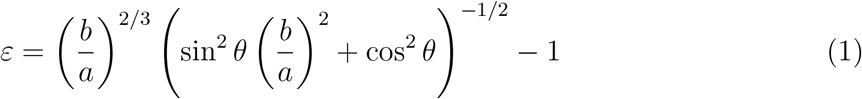

A detailed derivation of Eq. 1 can be found in Supplemental Note 1. Since the droplets were strained uniaxially, we obtained the effective tensile creep compliance along an arbitrary direction as

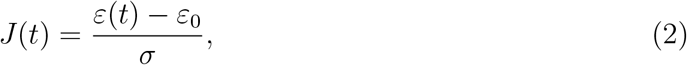

where *ε*_0_ is the average strain before the application of the magnetic field and *σ* is the magnitude of the applied magnetic stress introduced at *t* = 0.

To obtain a rheological description of the retinal tissue relevant to the developmental timescales, we characterized the creep compliance *J(t)* over durations of up to one hour. To study material properties across the early stages of retina development, we measured *J*(*t*) using organoids from three independent experiment rounds and on days in vitro (DIV) ranging from 16 to 24. Within this time period, retina progenitor cells differentiate into the major cell types, including ganglion cells, horizontal cells, cones, amacrine cells, rods, and bipolar cells[19]. The retina becomes laminated, organizing the different neurons into distinct layers. DIV 20 corresponds to the day of birth of the mouse[19].

### B. Retina tissue remodels in a scale-free manner

We applied stresses and measured the compliance for a duration of up to 1 hour with a temporal resolution of 1.5 seconds (Fig. 1g). One intriguing finding is that, when forces are applied within the retina tissue, the creep compliance shows a scale-free behaviour across four orders of magnitude. We found this power law scaling with respect to time for all our measurements across organoid days and rounds (Fig. 2a). The quality of the power law fit is excellent, with *R*^2^ values above 0.95 for 15 of 16 measurements (Fig. 2a, Methods). To model such a system using a simplistic mechanical circuit we used a single fractional-order linear viscoelastic element called a springpot[33]. The springpot is fully described by just two parameters, *β* and *c*_*β*_, and provides the creep compliance as[34]

**FIG. 2:**
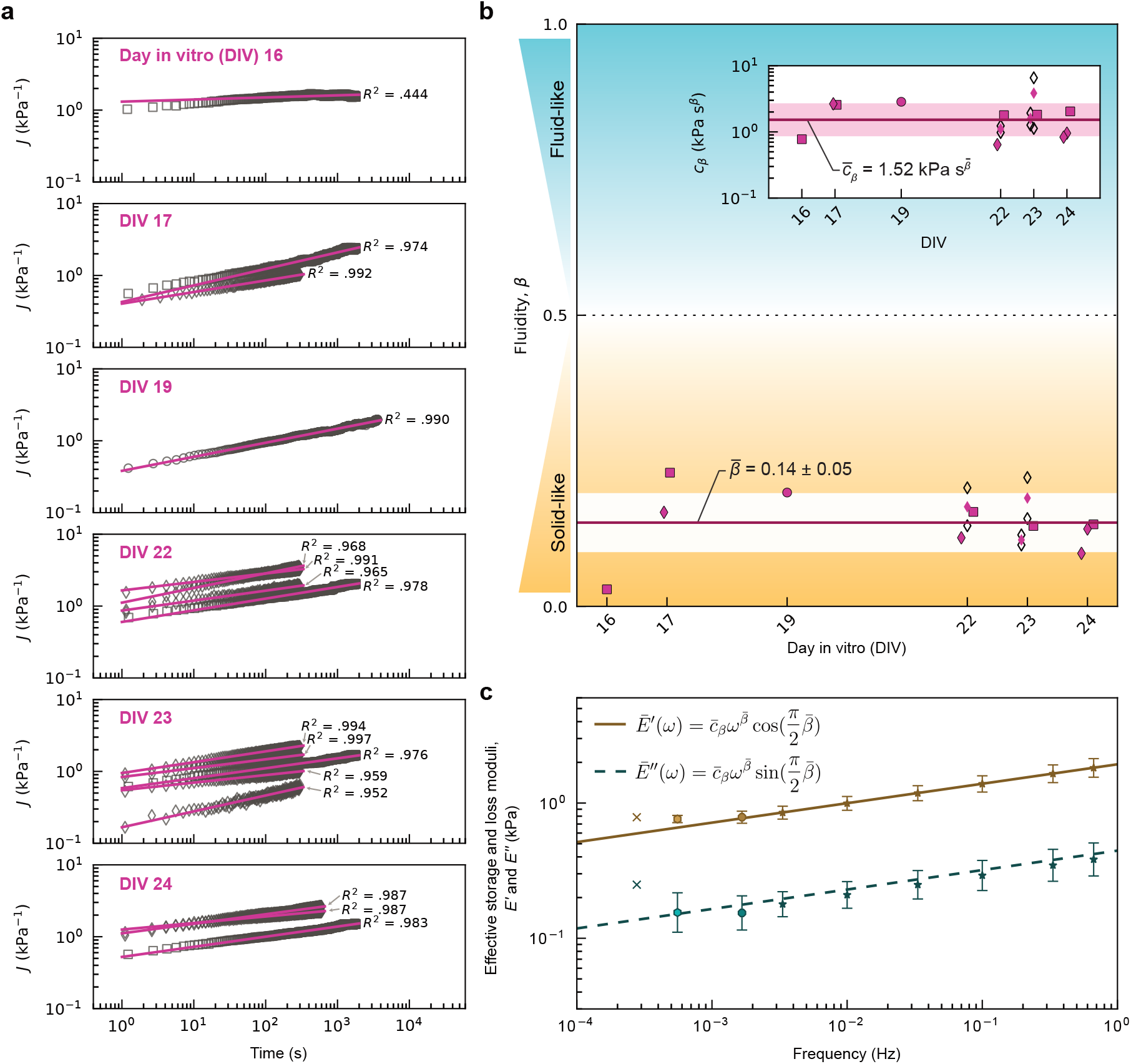
Scale invariance of strain relaxation in developing mammalian retina. **a**, Creep compliance in the retina tissue of mouse organoids ranging from DIV 16 to 24 are plotted against time. Lines show power law fits to data using equation 3 (Methods). **b**, The scaling exponents *β* obtained from (a) plotted against DIV remain smaller than 0.5, indicating solid-like behaviour. Inset: *c*_*β*_ plotted against DIV indicating the magnitude of viscoelastic resistance to the actuation. In **a** and **b**, measurements from different rounds of culture are plotted with different shapes (diamond, square, or circle). In the diamond round, pairs of compliance measurements were taken for organoids on DIV 22 and 23. These paired measurements are plotted in empty markers, and the average of these values is plotted in a filled marker. Lines indicate average values across all organoids, and the bands indicate *±* one standard deviation of the mean. For 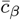, the geometric mean and standard deviation were used. **c**, The effective tensile storage modulus 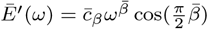 (solid line) and loss modulus 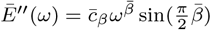 (dashed line) extrapolated across a range of frequencies. For select frequencies, the geometric average storage and loss moduli are plotted. The marker shape (x, *n* = 1; hexagon, *n* = 6; octogon, *n* = 8; star, *n* = 13.) indicates the number of organoids *n* represented in the data point. Each geometric average computed at frequency *f* reflects all measurements in which the maximal observation time *t*_max_ *≥* 1*/f* . Error bars indicate geometric standard error of the mean.

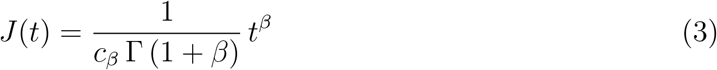

where Γ() is the Gamma-function. The dimensionless scaling exponent *β* tunes the response of the element from purely elastic, *β* = 0, to purely fluid, *β* = 1, with intermediate values describing the relative fluidity of the viscoelastic response. The parameter *c*_*β*_ has units of Pa*·*s^*β*^, reflecting the magnitude of viscoelastic resistance to displacement and flow. From fittings to the compliance data (Methods), we found 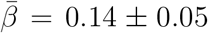 (mean *±* s.d.) and 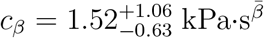 (geometric mean, geometric s.d.). These values for *β* demonstrate that the retina tissue, while still viscoelastic, is more solid-like than fluid-like (Fig. 2b).

The springpot model is an idealized power law material. Within this model, the storage modulus (*E*^*′*^) and loss modulus (*E*^*′′*^) are each described by single power laws[33], namely:

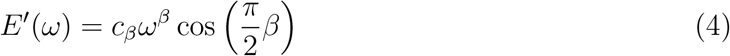

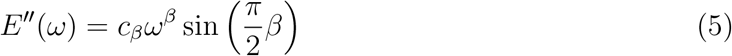

We adopt the springpot framework to calculate the effective tensile storage and loss modulus of the tissue across a range of frequencies using the experimentally determined values of *β* and *c*_*β*_ (Fig. 2c). Since *β <* 0.5, the storage modulus always dominates over the loss modulus (Fig. 2c), highlighting the elastic behaviour. The scaling behaviour *E*^*′*^, *E*^*′′*^ *∝ ω*^*β*^ together with the magnitude of *β* are consistent with the properties of a soft glassy material above the glass transition as discussed by Sollich and coauthors[35].

### C. Quantifying mechanics on the timescales of retinogenesis

Our measurements in organoids revealed the time dependence of the elastic and viscous characteristics of retinal tissue during development. Over timescales of 10^4^ *−* 10^5^ seconds, events like interkinetic nuclear migration, cell division, and differentiation collectively establish the architecture of the neural retina (Fig. 3a)[22, 23]. Using the measured values of *β* and *c*_*β*_ from the droplet actuation experiments in organoid tissue (Fig. 2), we determined the tissues’ effective mechanical properties on these developmentally relevant timescales. We found that the effective Young’s modulus *E* = (5*/*12)*E*^*′*^ decreased following a weak scaling (*E ∝ τ*^*−*0.14^) for timescales *τ* ranging from 1 second to 10^4^ seconds (Fig. 3b). On 10^4^ second timescale, the organoids become significantly softer with *E* = (0.23 *±* 0.07) kPa (Fig. 3c). Intriguingly, our measurements show that the relative variation in stiffness between different organoids is reduced. This is seen in the decrease in the coefficient of variation (CV=std/mean) from 1 to 10^4^ seconds (Fig. 3d). This suggests that stiffness variation becomes more tightly regulated at longer timescales such that initial heterogeneity between biological sample is reduced.

**FIG. 3:**
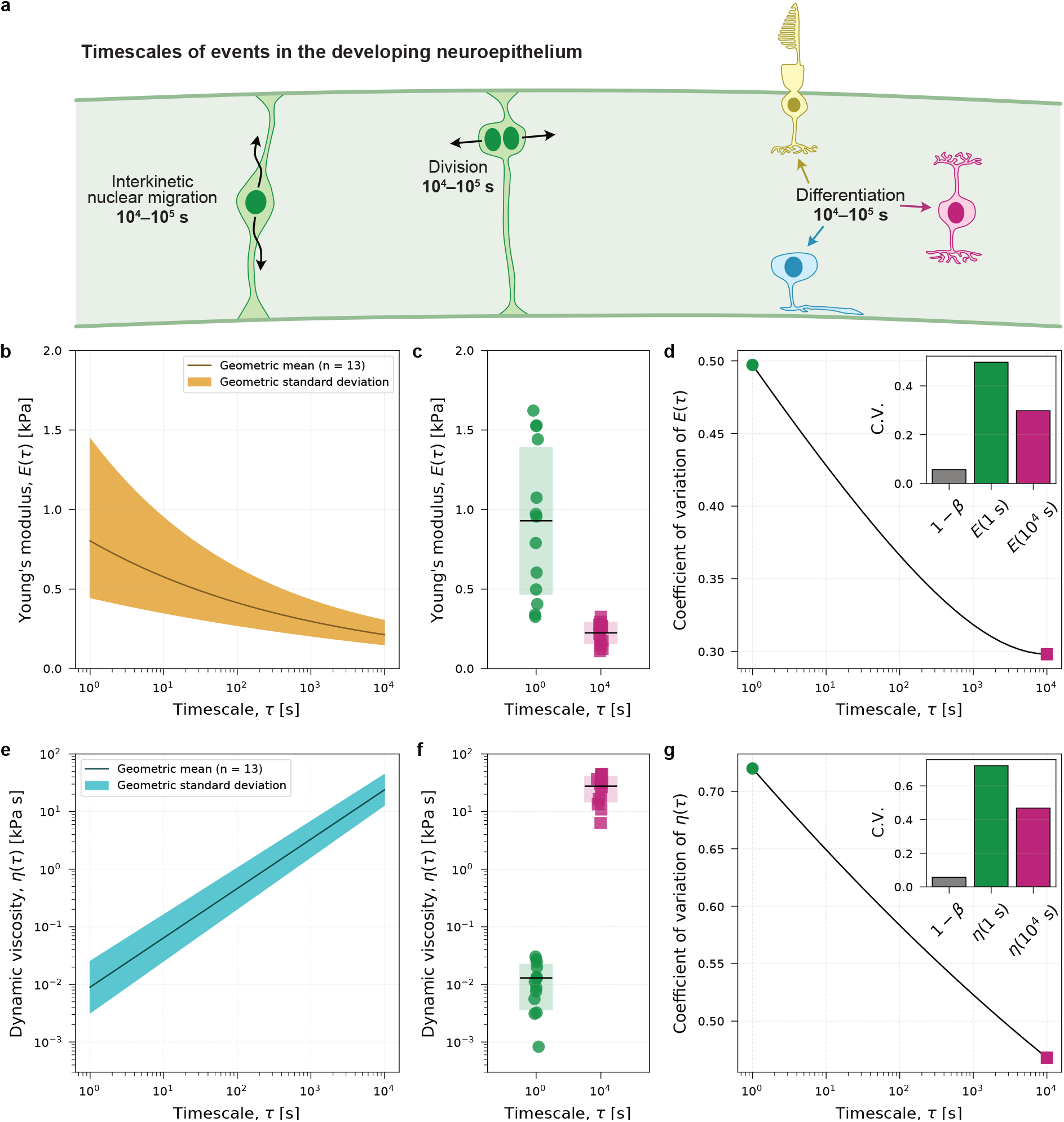
Quantifying the mechanics of the cellular mircoenvironment during retina development. **a**, The development of neuroepithelial tissues, like the retina, requires force-driven nuclear and cellular rearrangements to transform from a pseudo-stratified epithelium into a laminated structure of different cell types[22, 23]. **a**, Young’s modulus 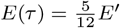 as a function of timescale *τ* (*n* = 13 organoids). **b**, Young’s modulus evaluated for each organoid at *τ* = 1 s and 10^4^ s. Lines and boxes indicate mean and standard deviations. **c**, Coefficient of Variation, CV(*E*) = s.d.(*E*)/mean(*E*) evaluated from *τ* = 1 s to *τ* = 10^4^ s, with inset comparing select values to CV(1 *− β*). **e**, Dynamic viscosity 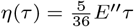 as a function of timescale *τ* (*n* = 13 organoids). **f**, Dynamic viscosity evaluated for each organoid at *τ* = 1 s and 10^4^ s. Lines and boxes indicate mean and standard deviations. **g**, Coefficient of Variation, CV(*η*) = s.d.(*η*)/mean(*η*) evaluated from *τ* = 1 s to *τ* = 10^4^ s, with inset comparing select values to CV(1 *− β*). **(b**,**e)** Lines indicate geometric means *E*_GM_ and *η*_GM_. Bands indicate geometric standard deviation.

We also determined the dynamic viscosity *η*(*τ*) = (5*/*36)*E*^*′′*^*τ* from the measured values of *β* and *c*_*β*_. For timescales ranging from 1 to 10^4^ seconds, we see a strong timescale dependence (*η ∝ τ* ^0.86^; Fig. 3e).

We found that the viscosity increases over three orders of magnitude from 13.0 *±* 9.3 Pa*·*s (mean *±* s.d.) at 1 second to 28 *±* 13 kPa*·*s at 10^4^ seconds(Fig. 3f). We expect that this increase in effective viscosity on longer timescales affects the timing and efficiency of neurons reaching their destination in the tissue as they navigate through this microenvironment. On these longer timescales, we observed a decrease in variability in the viscosity between different organoids, similar to that observed for stiffness (Fig. 3d).

We note that, in contrast to effective viscosity and elasticity, which are timescale dependent in both mean and coefficient of variation, the solidity (1 *− β* = 0.86 *±* 0.05) is a timescale independent metric.

Mechanical investigations in cells show that they increase their stiffness by becoming more solidlike (i.e. decreasing *β*)[36]. To investigate whether solidity and stiffness are coupled in the same way in the retina, we measured how stiffness varied across days in vitro and against different values of solidity. Like *β*, stiffness did not trend across different days of culture (Suppl. Fig. 5a,b). Interestingly, organoids with higher stiffness on short timescales were found to be less solid-like (Suppl. Fig. 5c), in contrast to the expected relation in cell mechanics. On the longer timescale of 10^4^ seconds, stiffness and solidity were not correlated (Suppl. Fig. 5d). These results highlight how solidity and stiffness are distinct material descriptors which can be tuned independently in tissues but are coupled on a cellular scale.

## III. DISCUSSION

The electrically functional neuronal architectures of the retina and brain are built by cells forcing their way through dense microenvironments. Neurons are subject to the rheological responses at the timescales of tissue development. Performing rheological probing on these timescales is difficult experimentally, but critically important for understanding the mechanics of cell movements and the mechanical cues driving differentiation. The main hurdles are (I) applying hour-long forces (II) within the microenvironments in which cells undergo neurogenesis (III) in mammalian neuronal tissue. Here, we overcome these challenges and provide a rheological description of the microenvironment of developing mammalian nervous tissue on second to hour timescales.

Our measurements revealed the previously undescribed rheological environment of neurogenesis (Fig. 1). This allows us to tackle key questions: what are the mechanical cues cells sense as they differentiate and what are the physical challenges of moving through this environment? On timescales ranging from 1 to 10^4^ seconds we discovered scale-free tissue remodeling (Fig. 2). Using a minimalistic theoretical model[33], we quantified the effective Young’s modulus and the dynamic viscosity experienced by neurons and neural progenitors across these timescales (Fig. 3). Throughout the observation window of 16 – 24 days *in vitro*, corresponding to the *in vivo* window of embryonic day 16 to postnatal day 4, we found neuronal tissue remodeling is well described by a power law model with an average exponent of 0.14, indicating solid-like behavior during this period of significant cell maturation and tissue transformation.

Our findings provide an experimentally grounded starting point for understanding the biophysical challenges of dynamically self-organizing a functional neuronal network. Traditionally, tissue mechanics are described by reporting values for Young’s modulus or dynamic viscosity characterized on second timescales or faster[4, 11]. Using ferrofluid droplets as mechanical actuators we show the significance of time-dependence in the rheology of the forming retina. From one to ten thousand seconds, the effective Young’s modulus decreases fourfold, while the effective dynamic viscosity increases by over three orders of magnitude. Our measurements of retina Young’s modulus determined at 1 second (0.93*±*0.46 kPa; mean *±* s.d.; Fig. 3c) compare very well with values of bulk modulus previously obtained from guinea pig and mouse retina via AFM[7, 10]. Our measurements recapitulate prior characterizations of retina mechanics at seconds timescale, but now offer a significant advance: a rheology describing remodeling mechanics reaching into the regime of developmental events.

Our discovery of scale-free rheology provides a minimalistic biophysical description of how developing neuronal tissues respond to active cellular forces and what mechanical cues these cells in turn sense. In vivo studies in Xenopus show that axon growth is regulated by environmental stiffness, with a Young’s modulus around 0.6 kPa characterized at seconds timescale[2]. In contrast, cell culture studies indicate that synapse formation occurs over minutes to hours[37] highlighting the need for measurements and theoretical models linking these timescales. The fractional-order model we implement here combined with experimental characterization bridges this gap and advances mechanobiology by clarifying the role of time-dependent material properties in neurogenesis[23]. Seminal cell culture experiments have shown that key cell behaviours, including differentiation, are regulated via material properties such as the elastic modulus and viscoelastic properties of the cellular microenvironment[38, 39]. While cells’ ability to sense mechanical cues is well established, understanding the nature of this mechanotransduction is still an active area of research[39– 41]. Here, by characterizing the time-dependent tissue rheology, we provide a framework for understanding the mechanical signals sensed by cells and for tackling open questions about how time-dependent material properties affect neuronal network formation.

The scaling behavior we observed indicates that soft glassy rheology (SGR) could be a framework for understanding retinal development from a soft matter physics perspective. In their seminal theoretical paper, Sollich et al[35] explore the rheology of so-called soft glassy materials under different levels of effective temperature or noise. In the low activity regime, particles comprising this material are “caged” or confined to their local environment by a landscape of energy wells. Increasing activity in the system tunes the rheology through a glass transition from solid-like to fluid-like by increasing the rate at which particles “uncage” by hopping out of their energy wells . In our measurements, the scaling of the storage and loss modulus as *ω*^*β*^ with *β* = 0.14 is consistent with such a soft glassy material near and above the glass transition[35, 42, 43]. Indications of SGR have been seen previously in 2D cell culture[44] and in 3D in non-neuronal tissues[43, 45]. However, the role of soft glassy physics for the development of neuronal systems is yet to be explored. SGR offers an exciting picture for neurodevelopment in which the proximitiy from the glass transition is tuned through interactions between cells which determines the height of the energy wells, and through cellular activity which determines the effective temperature[46]. Therefore, we hypothesize that a glassy material just above its glass transition could be one way to balance the competing engineering goals of retina development: maximizing retina imaging resolution requires a dense, regular packing of neurons; at the same time, tissue growth necessitates the dynamic refinement of the architecture. While such a hypothesis demands more evidence than similarity in the scaling parameter, the ability to directly probe tissue rheology through local mechanical actuation with magnetic droplets puts us in a position to test ideas from SGR, like stress-induced uncaging behaviour.

Power law rheology has been observed in single cells at shorter timescales, and our scalefree observations in tissues quantitatively align with findings in 2D cell culture[42, 47, 48]. Magnetic tweezer experiments on the cytoskeleton of various cell lines found *β* between 0.17 and 0.25[42, 47, 49], comparable with what we report here at completely different length and time scales in 3D neuronal tissue. In cell culture experiments, cells with higher stiffness (higher elastic modulus) are typically found to be more solid-like (lower scaling exponent)[36, 42]. In contrast, we found that the stiffest organoids were those with the highest scaling exponents (Suppl. Fig. 5c). This observation suggests that stiffness and the scaling exponent may be independently regulated, and cautions against interpretation of a decrease in stiffness as an indication of increased tissue fluidity. We observe a surprisingly small spread in the measurements of the scaling exponent *β* (Fig. 2b). This lets us speculate that *β* is a tightly regulated physical property of the tissue. Intriguingly, we found the relative spread in elastic modulus across biological replicates to significantly decrease for longer timescales. The tuning of variances of biochemical factors[50] has been shown to play a key role for development. Our results trigger the question what role the mechanical variances play and how those control neurogenesis. Whether this behaviour is reminiscent only of retina tissue or a general hallmark of other neuronal systems is an open question which could be explored by applying the concepts presented here in other organoid systems.

Direct experimental measurements are critical for elucidating how physical properties of the tissue emerge from interactions at the cellular level. In this context, our work informs recent theoretical vertex models with experimentally measured parameters. Notably, vertex models successfully capture a jamming-unjamming transition in epithelial sheets[46]. A jamming transition has been shown experimentally in non-neuronal tissues[26, 51, 52], and inferred in the non-mammalian retina indirectly from imaging of tissue architecture[53]. Here, we provide the first direct quantification of neuronal tissue mechanics over timescales from seconds to hours, uncovering a scaling of both storage and loss modulus with *ω*^*β*^ that challenges predictions from recent 3D vertex models[54, 55].

This work introduces three significant advances in the use of magnetic drops for material characterization: first, an extended the duration of applied stress up to one hour; second, a radically simplified calibration method which is additionally validated in viscoelastic hydrogels; and third, an incorporation of fractional rheological models into the analysis and interpretation of magnetic droplet-based measurements in tissues which was previously lacking.

This discovery of scale-free rheology in neuronal tissue brings concepts developed in the soft matter physics community into the purview of developmental and neurobiologists. While the electrical and informational properties of the nervous system have been the subject of much investigation, the physical material properties critical to the self-organization and function have not been adequately explored. The description of neuronal mechanics as introduced here could be the key to unlocking its role in processes ranging from migration and proliferation, to cortex folding and synapse formation, ultimately linking mechanics, network architecture and functionality[1].

## IV. METHODS

### Stem cell culture and retina organoid generation

We generated organoids from mouse embryonic stem cells in three independent rounds. For each round, we thawed Rx-GFP K/I EB5 cells (AES0145, RIKEN) from LN2 storage at passages 26, 23, and 23, respectively, and maintained them for two passages according to a previously published protocol[16, 29] prior to retina differentiation. We seeded 3 *×* 10^3^ cells per well in 100 µl volumes of Retinal Differentiation Media (RDM) in 96-well ultra low adherent round bottom plates on DIV 0 and added 2% Matrigel on DIV 1. Cells and organoids were cultured at 37°C. We cultured organoids at 20% O_2_ until DIV 7, before transferring to Petri dishes for further culture in retina maturation medium 1 (RMM1) at 40% O_2_. On DIV 10, we transferred organoids to new dishes containing retina maturation media 2 (RMM2) freshly supplemented with all-trans retinoic acid (0.5 µM) and taurine (1 mM). On DIV 14, we screened retina organoids for GFP under a Nikon SMZ25 stereoscope with a 0.5X SHR Plan Apo objective and fluorescence illumination and transferred GFP-positive organoids intact (without trisection) to new dishes with RMM2 freshly supplemented with taurine (1 mM). We maintained organoids up to DIV 26 in RMM2 freshly supplemented with 1 mM taurine, removing and replacing half of the media on DIV 16, 18, 21, 23, and 25.

### Immunohistochemistry

A selection of organoids from the first round of culture in which we made mechanical measurements were fixed in 4% PFA for 60 minutes on an orbital shaker at 4°C. Following fixation, organoids were rinsed three times in PBS. After four days in PBS at 4°C, organoids were transferred to 30% sucrose and held overnight at 4°C, before being embedding and frozen in optimal cutting temperature compound. Organoids were cut into 10-30 micron thick sections on a cryotome (CryoStar NX70). Sections were held at 4°C prior to staining. We performed immunohistochemistry using recoverin Rabbit IgG (1:500) and goat anti-rabbit AF568(1:500)(Merck Millipore AB5585). We blocked sections (90% PBST; T = TritonX 0.5% and 10% NGS) for 1 hour at room temperature. Following a PBS rinse, we incubated sections in primary antibody overnight at 4°C and in secondary antibody for 4 hours at room temperature, with PBS rinses between and afterwards. Finally, we stained sections with Nucspot650 (Biotium) (1:500) for 30 minutes before a final wash. We mounted sections in FlourmountG on a glass cover slip and stored samples at 4°C.

### Preparation and injection of ferrofluid

We prepared magnetic oil for injection experiments from a 1:1 (v/v) mixture of 0.2 micron filtered DFF2 (Ferrotec) and Fluorinert FC40 (FL-005-HP-1000, 3M) with 2.5 wt.% 008-FluoroSurfactant (RAN Biotechnologies) and 250 µM F86Cy5[31]. To control the concentration of DFF2 across multiple rounds and to compensate evaporation, we stored the magnetic oil at 4°C and thinned it with Fluorinert FC40 as needed. Prior to injection, we sonicated the magnetic oil for 10 minutes at room temperature before loading it into a 55 mm long straight spiked 10 micron ID ICSI-Pipette (BM100T-10P, BioMedical Instruments).

We injected organoids with ferrofluid droplets under a Nikon SMZ25 stereoscope (see above) on agarose under RMM2 media. Immediately after organoid injection, we injected ferrofluid into N190000 viscous standards (Cannon Instrument Company) to calibrate the applied stress (Suppl. Information).

### Imaging of living and fixed organoids

Prior to injection, we acquired fluorescence and phase contrast images in Fig. 1a using a Ximea xiC camera mounted to a Zeiss Axiovert 200M inverted fluorescence microscope with condenser. After droplet injection, we maintained organoids at 37°C in a humidified 5% CO_2_ 40% O_2_ environment for approximately 4 hours. For live imaging, we placed organoids within a silicone well sandwiched between two glass coverslips and filled with supplemented RMM2 before transferring to the 37°C live imaging chamber (Digital Pixel). We performed all confocal microscopy on a Zeiss LSM 980. We imaged the fixed section in Fig. 1b with a C-Achroplan 32x/0.85 water immersion objective. We imaged droplets in stress calibration measurements in viscous standards with the A-Plan 10x/0.25 air objective and the droplets for mechanical measurements in organoids with either the Plan-Apochromat 20x/0.8 M27 air objective or the LD LCI Plan-Apochromat 40x/1.2 Imm Korr DIC M27 objective. We acquired all time series at a rate of 1 frame per 1.5 seconds and frame scan time of 0.2 seconds. To improve visibility for the reader in Fig. 1e, we used Fiji to sum images from a mechanical measurement time series (DIV 19, Fig. 1f,g and Fig. 2a) across 10 frames.

### Application of magnetic stress

We applied a step change in stress to the magnetic droplet by rapidly moving a Hallbach array of permanent magnets mounted to a LabVIEW controlled linear stage, as previously described[25], from an initial position far from the droplet (low stress, *σ*_*m*_ = *σ*_0_) to a final position close to the droplet (high stress, *σ*_*m*_ = *σ*_0_ + *σ*), while imaging the droplet with confocal microscopy. In order to determine the orientation, timing, and magnitude of the applied stress in the organoid, we applied the *same* magnetic field to droplets in viscous standards (N190000, Cannon Instrument Company) at (37 *±* 0.1)°C.

### Calculation of strain

In the absence of other stresses, the droplet strain under a uniaxial stress is *ε* = (*b/a*)^2*/*3^ *−* 1. When other stresses may be present in the tissue, the droplet shows a non-zero initial deformation. To calculate the strain in the direction of the applied field in the presence of pre-deformations we use the droplet’s aspect ratio, *b/a*, and declination, *θ*, relative to the magnetic field (Figure 1e). We measured the magnetic field orientation from the alignment of the elliptical segmentation of the droplet at maximal strain in calibration measurements. Following the example of [32], it can be shown that the angle-corrected strain along an arbitrary axis is given by 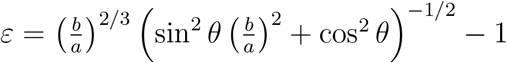. A full derivation can be found in Suppl. Note 1.

### A python-based user-friendly interface for droplet shape analysis and fitting

To be able to perform calibrations, image processing and analysis efficiently on long time series we developed a python-based user interface using libraries like NumPy, SciPy, and OpenCV. The pipeline consists of three main tabs – analyzing, calibrating, and fitting. For a screenshot, see Suppl. Fig. 1. The graphical user interface (GUI) allows users to fit ellipses to droplet images from time-lapse sequences. This process generates strain curves, which are used in a next step to extract the material’s mechanical properties. The software is programmed in a modular way using separate classes for the GUI, image analysis, and fitting algorithms. The fitting algorithm uses the curve_fit function from SciPy[56] to model pre-magnetic and the magnetic field phase (creep). The best fit is selected based on minimal residuals by iterating 100 times over statistically varied initial conditions. The pipeline also features error propagation using the uncertainties[57] package, providing uncertainty estimations for fit parameters. A calibration measurement routine ensures the accuracy of the applied stress by accounting for experimental errors, including fitting errors and errors in the measurement of the viscous standards.

### Modeling organoid mechanics

We modelled the mechanical response of the droplet in the organoid as a single springpot element characterized by the two parameters *β* and *c*_*β*_ giving the creep compliance 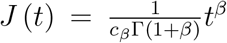. We obtained *β* and *c*_*β*_ from a linear fit to the log transformed time and compliance data. In contrast to the Kelvin-Voigt model used for calibration measurements, our model for the strain dynamics in the organoid does not consider the droplet’s capillary forces as a parallel spring element, as we could not observe any stiffening effect on the long term behaviour of the creep compliance. See Suppl. Fig. 3 for a comparison between the single springpot model and a springpot in parallel with a spring. While other models were considered (see Suppl. Fig. 4), the springpot is the simplest model describing the data across several orders of magnitude. We obtained values for the Young’s modulus and dynamic modulus from the parameters of the springpot model (*β* and *c*_*β*_) following the example of [33], and applying numerical factors to relate the 1D tensile storage and loss modulus to the 3D Young’s modulus and dynamic viscosity, as was done previously[25].

### Calibration of applied stress

To determine precisely how much stress was applied during a measurement in organoid tissue, for each day of organoid measurements we injected multiple droplets (typical diameters (85 *±* 35) microns) of the same magnetic oil used in the organoid into a viscous standard (N190000, Lot no. 14101d, Cannon Instrument Company). To control the viscosity, the imaging chamber and sample were held at (37.0 *±* 0.1)°C while calibration experiments were performed. We acquired fluorescence confocal time series of the droplet (1.5 second interval time) while introducing a magnetic stress, as described above, in the same manner as in the organoid. From these image series, we obtained strain versus time curves using our custom python-based GUI. We determined the magnitude of stress change *σ* and the time *t*_0_ corresponding to the change by modelling the droplet in viscous standard as a 1D Kelvin-Voigt material with spring *k*_*d*_ and dashpot *µ*, as before[25]. In this 1D description, the magnetic stress reads *σ*_m_ = *k*_d_*ε* where *ε* is the equilibrium droplet strain and the droplet spring constant 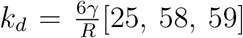. Under a step change in stress from *σ*_0_ to *σ*_0_ + *σ*, we model the strain dynamics as 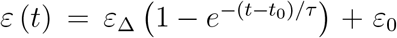, where *t*_0_ is when the step change occurs, *ε*_0_ is the initial strain before *t*_0_, *ε*_Δ_ + *ε*_0_ is the final strain at equilibrium, and *τ* = 6*ηR/*5*γ* is the relaxation timescale set by the surrounding medium’s viscosity (*η*), the droplet radius (*R*), and the interfacial tension (*γ*)[60]. The timescale in our 1D model *τ* = *µ/k*_d_ gives us the relationship between the dashpot *µ* and dynamic viscosity *η* as *µ* = (36*/*5)*η*. We obtained values for *τ*, *t*_0_, and *ε*_Δ_ by directly fitting the strain versus time data, and we measured the viscosity of N190000 at 37°C to be *η*_*R*_ = (214.5 *±* 6.8) Pa*·*s (mean *±* s.e.m.) using a rheometer (see Methods below), allowing us to measure *k*_d_ and *σ*.

Compared to the rheometer data, the droplet technique shows very good agreement for materials with viscosities higher than those used for calibration; for example, testing with the Cannon Instrument Company N2700000 viscosity standard results in a relative error 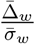 of 4.3% (see Suppl. Fig. 2), where 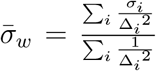 is the weighted mean stress and 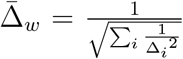 is the weighted standard deviation. The total error of the applied stress 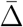 contains also the systematic error of the viscosity standard using 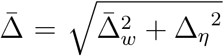, where 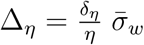. When calibrating with N190000 at 37°C, the uncertainty in the viscosity from the rheometer measurement (*δ*_*η*_*/η* = 3.2%) is usually the largest source of error. When considering uncertainties from the fit as well, a typical total relative error in the applied stress amounts to 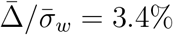.

### Rheometer measurements of viscosity standards

The dynamic viscosity *η*_*R*_ of the viscous standards (Cannon Instrument Company) was determined using a shear plate rheometer (Kinexus Ultra+, Malvern Instruments Limited). We first performed frequency and amplitude sweeps to determine the range of frequencies and strains, respectively, where the viscosity remained constant. Then we performed the measurements at a constant frequency and strain that were 10 % smaller than the upper limits of these ranges. For each viscous standard we performed 3–5 measurements at 37°C that lasted between 485–515 seconds. Of each data set, the last data point was dropped and the remaining last 20 data points were averaged to obtain *η*_*R*_. To test that this method yields accurate results of the standards’ viscosities, we also probed the standards N62000 and N190000 at 25°C to compare the results with the values given by the standards’ manufacturer for the lots used (Suppl. Table 1).

### Preparation of collagen gels

All collagen gel samples were prepared to final concentrations of 1.76 mg/mL following a standard protocol[61]. Collagen samples were prepared in final volumes of 300 µL by first combining 30 µL of 10x PBS, 3 µL of 1 N NaOH and 117 µL of diH20, and then mixing in 150 µL of type I tail collagen (Corning 354236, 3.52 mg / ml). To prevent premature gelation, all mixing steps were performed on ice.

### Rheometer measurements of collagen gel

Rheological measurements of collagen gels were performed with a shear rheometer (Kinexus Ultra+, Malvern Panalytical). To prevent slip, Poly-L-Lysine-coated 25 mm glass cover glasses were glued to the upper and lower plates of the rheometer using a cyanoacrylate adhesive (PATTEX) following previously established protocols [62].

The sample was loaded into the rheometer at a temperature of 25°C. A 210 µL volume of collagen solution was transferred to the lower geometry and the upper geometry was then immediately lowered to a gap of 0.4 mm. Mineral oil (Sigma-Aldrich) was applied to the exposed boundary to prevent evaporation.

The temperature of the rheometer was increased to 37°C to initiate gelation. During this process, a single-frequency oscillation test was performed at a shear strain amplitude of 0.01% and a frequency of 0.159 Hz (1 rad/s). This allowed monitoring of the development of the shear storage modulus (*G*^*′*^) and the loss modulus (*G*^*′′*^) over time. Once the loss modulus reached equilibrium, the creep test was started.

Creep tests were performed by applying a shear stress of *σ* = 10 Pa and measuring the shear strain for at least 200 seconds with a time resolution of at least 1 second. A new sample was used for each measurement. Measurements in which slip was detected were excluded.

The values for scaling exponent *β* were determined by fitting power laws to the shear strain versus time data, considering only data collected between 1 and 200 seconds from the start of the shear stress. The values for *c*_*β*_ were obtained by equating equation 3 to the shear creep compliance as defined in equation 4:

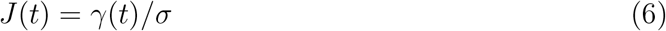

The Young’s modulus *E* was determined from the shear modulus *G* by assuming a Poisson’s ratio *v* = 0.5. The shear modulus was calculated for the timescales 2*π/ω* using equation 5: [33]

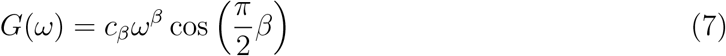

### Magnetic droplet-based measurements of collagen gels

For each measurement, the collagen solution was transferred immediately after mixing to a prefabricated silicone well affixed to a cover glass. Working on ice, the ferrofluid was injected into the collagen to form droplets of diameters between 0.14 and 0.16 mm. The sample volume was then capped with a second cover glass to form a sealed imaging chamber. All samples were incubated for at least 30 minutes at 37°C prior to mechanical characterization.

Magnetic droplet creep tests were performed at 37°C as described above for three samples on three different days. The images were acquired with a frame time of 0.2 s, a timestep of 1.5 s, and a duration of at least 200 s. For a quantitative comparison of Young’s modulus between techniques, the applied stress was determined to be *σ* = 21.5 Pa by calibration measurements performed on one of the days.

The scaling exponent *β* was determined by fitting a power law to *ϵ − ϵ*_0_ versus time data, considering only data collected up to 200 seconds from the start of magnetic stress. The value for *c*_*β*_ was determined by equating equation 3 to equation 2, and the Young’s modulus *E*(*ω*) was determined from *β, c*_*β*_, as was done in the organoid measurements, using equation 6:

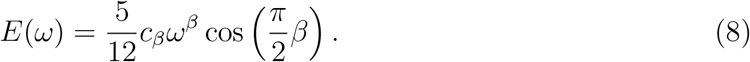

## Author contribution

E.R.S and F.S. conceived the research. R.J. cultured the mouse retina organoids. M.F., E.R.S and F.S. developed the analysis pipeline. E.R.S, M.F. and A.T.B. analyzed the data. E.R.S., M.F., and L.M.H. performed the droplet injection experiments and confocal imaging. A.T.B. and F.S. probed the viscous standards on the rheometer. R.J. performed the immunohistochemistry. L.M.H. probed the collagen gels with the rheometer. E.R.S. performed the statistical analysis. All authors contributed to the manuscript.

## ACKNOWLEDGMENTS

We thank Timo Betz and Max Bi for stimulating discussions. We thank Ellen Sletten for providing the fluorinated dye. We thank Mythili Padavu, Georgii Volkov, Teresa Rogler, Vlasta Hadalin and Lena Glanz for support with the organoid culture. We thank Max Bi and Marie Lackmann for feedback on the manuscript. We thank Jacopo di Russo, Florian Huhnke and Joachim Spatz for their contribution in the early phase of the project. We thank Joachim Rädler for his support. This work was supported by the European Research Council (ERC) under the European Union’s Horizon 2020 research and innovation program (Grant Agreement No. 850691), the Center for NanoScience (CeNS) at the Ludwig-Maximilians-University (LMU), Munich, and the Munich Cluster for Systems Neurology (SyNergy). The work was supported by the Human Frontier Science Program (Grant No. 18807).

