## Supplemental Information for "Scale Invariance of Mechanical Properties in the Developing Mammalian Retina"

June 2025

### 1 Supplementary Movies

#### 1.1 Supplementary Movie 1 (caption only)

**Supplementary Movie 1. 3D rendering of confocal microscopy image of immunofluorescence staining of cryosection**. Recoverin, expressed in photoreceptor and a subset of bipolar cells is labeled (day in vitro 25).

#### 1.2 Supplementary Movie 2 (caption only)

**Supplementary Movie 2. Timeseries of retinal tissue (day in vitro 19) undergoing application and removal of stress by a magnetic droplet.** The droplet is shown in magenta and the GFP in green. In order to increase the signal to noise for the viewer, each frame in this movie was produced by summing over 10 frames in the original confocal time series. Movie frame cycle time: 15 seconds. Original acquisition cycle time: 1.5 seconds. Cell line: Rx-GFP; passage: 25; day in vitro: 19.

### 2 Supplementary Figures

#### 2.1 Supplementary Fig. 1

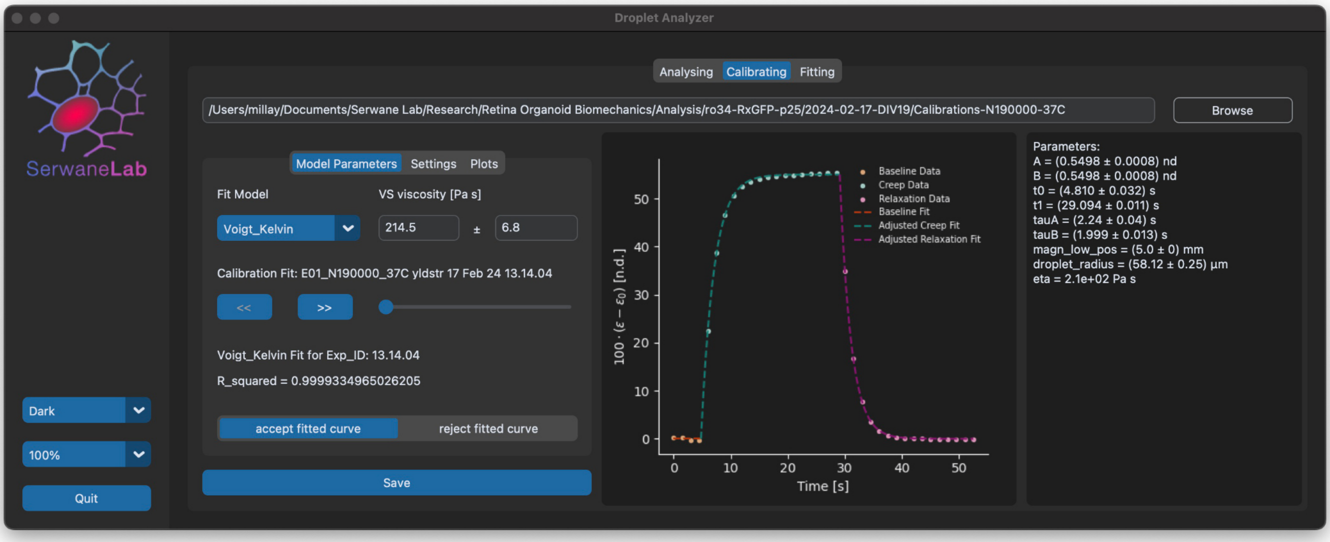

**Supplementary Fig. 1. Screenshot of the python-based GUI used for image and mechanical analysis.** Screenshot of the python-based GUI used for (1) obtaining strain versus time curves from image data and (2) fitting strain versus time curves. In this screenshot, we see the user calibrating the applied stress by fitting the strain versus time data from a measurement in the N190000 viscous standard (Cannon Instrument Company) with a Kelvin-Voigt model. At the time of publication, a version of this software can be found at `https://github.com/serwane/retina_mechanics`.

#### 2.2 Supplementary Fig. 2

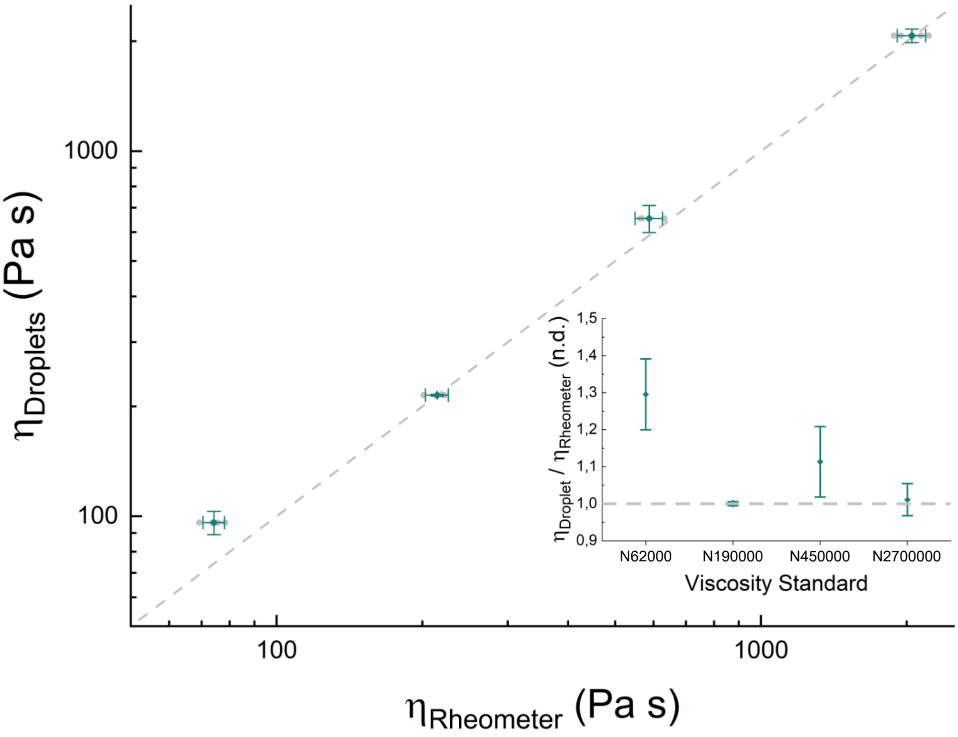

**Supplementary Fig. 2. Comparison of viscosity measurements between rheometer and magnetic droplet.** To validate the calibration of magnetic stress in the N190000 viscous standard (Cannon Instrument Company), we compared the viscosity $\eta_{\text{Rheometer}}$ measured with the shear plate rheometer (Kinexus Ultra+, Malvern Instruments Limited; See Suppl. Table 1 for the number of independently loaded samples for each standard, shown here in gray) with the viscosity $\eta_{\text{Droplets}}$ measured with magnetic droplets ($n = 19$ for N190000, $n = 12$ for N62000, N450000, and N2700000, where $n$ is the number of droplets injected into the viscous standards). Cannon Instrument Company high viscosity standards, from left to right: N62000, N190000, N450000, and N2700000. Error bars show ± one standard deviation.

#### 2.3 Supplementary Fig. 3

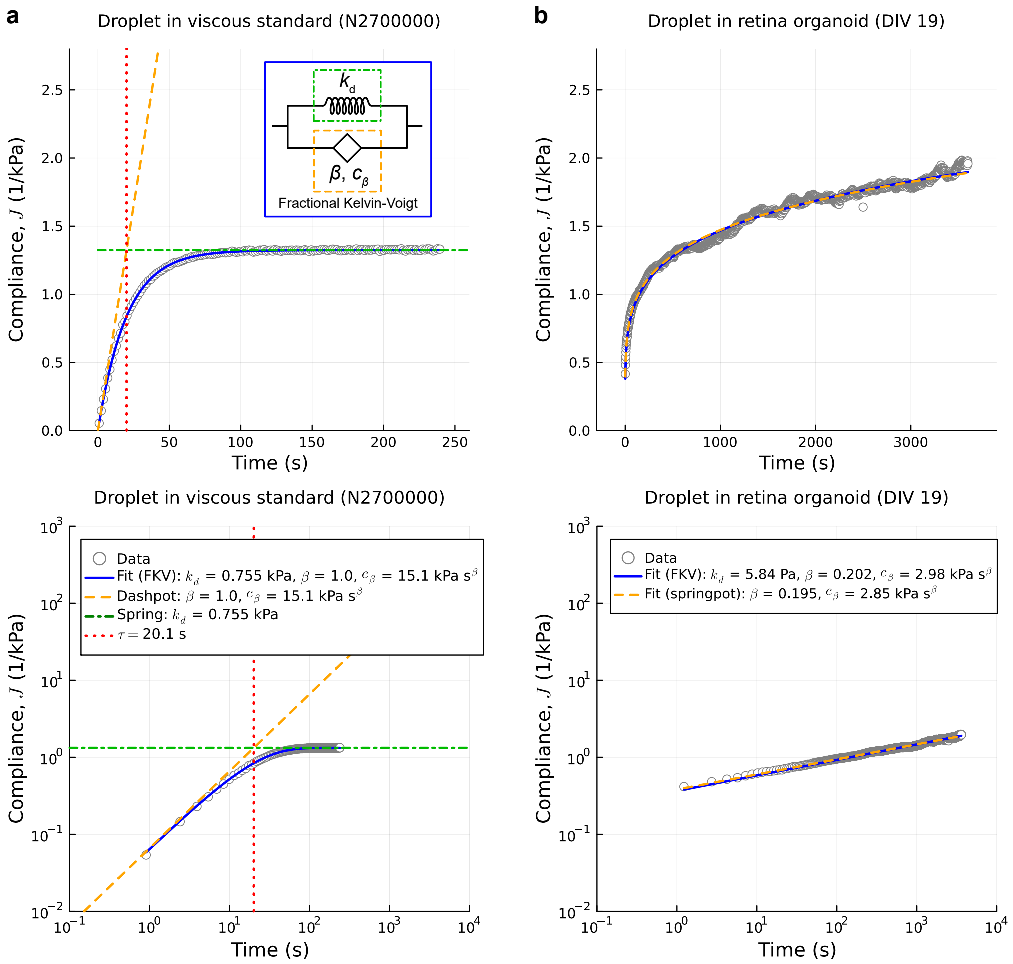

**Supplementary Fig. 3. Droplet interfacial tension (IFT) is not important to model measurements in retina organoid (RO). a,** Compliance versus time data for droplet in N2700000 viscosity standard at 37 °C (open gray circles; top axes, lin-lin; bottom axes, log-log). In the Fractional Kelvin-Voigt (FKV) model (inset, top), the spring ($k_d$) represents the droplet and the springpot ($\beta$, $c_\beta$) represents the sample, the viscosity standard in the dashpot limit $\beta = 1$, and $c_\beta = \mu = (36/5)\eta$. The FKV compliance is $J(t) = \frac{t^\beta}{c_\beta} E_{\beta,1+\beta}(-\frac{k_d}{c_\beta} t^\beta)$, where $E_{a,b}(x)$ is the generalized Mittag-Leffler function.[1] FKV fit (solid blue line), dashpot response (dashed red line), and spring response (dash-dotted green line) are shown. The crossover between regimes is at $\tau = \mu/k_d = 20.1$ s (dotted orange line). The droplet IFT in a viscosity standard must be modeled to describe the crossover from a short term fluid response to a long term solid response. **b**, Compliance versus time data for a droplet in RO (open gray circles; top axes, lin-lin; bottom axes, log-log). The similarity between FKV fit (solid blue) and springpot fit (dashed red) indicates short time regime. This is confirmed by the crossover time predicted by the fitted values: $\tau = (c_\beta/k_d)^{1/\beta} = 2.46 \times 10^{13}$ s. Since RO compliance data are clearly in the short time regime, we ignored the droplet IFT effects and fit power laws in our main analysis.

#### 2.4 Supplementary Fig. 4

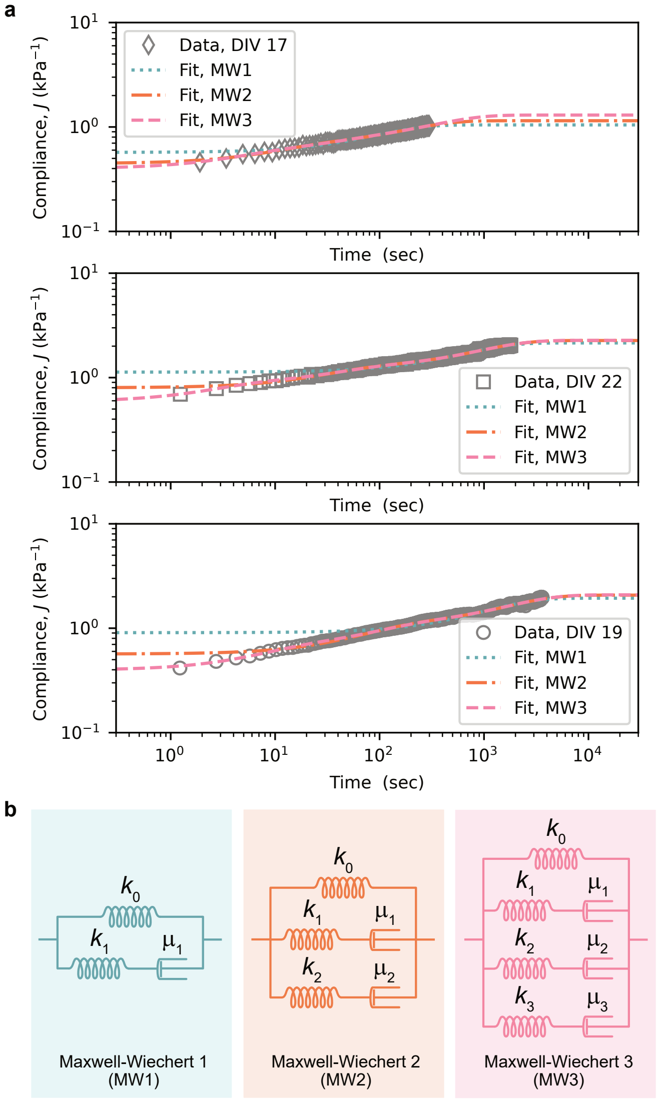

**Supplementary Fig. 4. Fitting compliance data using the Maxwell-Wiechert (MW) models.** **a**, Compliance data from 17 (top, diamond), 22 (middle, square), and 19 (bottom, circle) DIV retinal organoids, with 1, 2, and 3 branched Maxwell Wiechert (MW) model fits. These fits illustrate the challenge of fitting compliance data across more than three orders of magnitude with a simple model. Only the MW3, requiring seven total parameters, is able to provide a reasonable fit over the entire range. **b**, Schematics of the MW1, MW2 and MW3 models, showing the increasing complexity of the Maxwell-Wiechert models required to improve the range of prediction.

#### 2.5 Supplementary Fig. 5

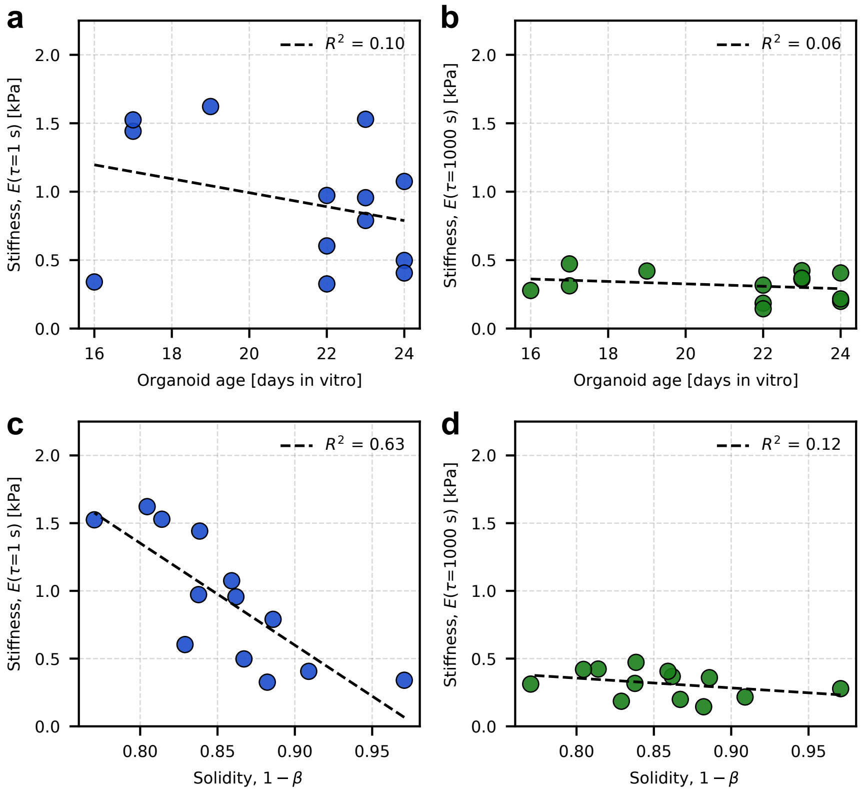

**Supplementary Fig. 5. Retina organoid tissue stiffness is more tightly dispersed on longer timescales.** **a**, Short-term organoid stiffness measurements (Young's moduli evaluated at 1 s) and **b**, Long-term organoid stiffness measurements (Young's moduli evaluated at 1000 s) plotted against organoid age.**c**, Young's moduli evaluated at 1 s plotted as a function of solidity $(1-\beta)$ and **d**, Young's moduli evaluated at 1000 s plotted as a function of solidity $(1-\beta)$ suggests that organoid tissue stiffness is anticorrelated with solidity on short timescales, but not long timescales. **(a-d)** Each data point represents measurement from one organoid. Dashed lines and $R^2$ values obtained from simple linear regression. $N = 13$ organoids.

##### 2.6 Supplementary Fig. 6

### 3 Supplementary Notes

#### 3.1 Supplementary Note 1

**Derivation of the formula for elliptical strain along arbitrary orientation**
To determine the strain of an elliptical section of an ellipsoid along an arbitrary direction, we can follow the example of Treagus[2]:

$$\lambda' = \frac{\lambda_1' + \lambda_2'}{2} - \frac{\lambda_2' - \lambda_1'}{2} \cos 2\theta \tag{1}$$

where $\theta$ is the angle relative to the first (long) axis of the ellipse. Let $b$ and $a$ be the long and short semi-axis lengths, $l$ be the length of the radial vector running from the center to the edge of the ellipse at a declination of $\theta$ from the long axis, and $R$ be the radius of a sphere with equal volume to the ellipsoid of revolution about the long axis of the ellipse. Then, $\lambda_1' = (R/b)^2$, $\lambda_2' = (R/a)^2$, and $\lambda' = (R/l)^2$. Let's now define the strain at orientation $\theta$ to be $\varepsilon \equiv \frac{l-R}{R}$ and find an expression in terms of $b$, $a$, and $\theta$.

$$\varepsilon = \frac{l}{R} - 1 \tag{2}$$

$$\varepsilon = (\lambda')^{-1/2} - 1 \tag{3}$$

$$\varepsilon = \left( \frac{\lambda_1' + \lambda_2'}{2} - \frac{\lambda_2' - \lambda_1'}{2} \cos 2\theta \right)^{-1/2} - 1 \tag{4}$$

$$\varepsilon = \left( \frac{(R/b)^2 + (R/a)^2}{2} - \frac{(R/a)^2 - (R/b)^2}{2} \cos 2\theta \right)^{-1/2} - 1 \tag{5}$$

$$\varepsilon = \left( \frac{(R/a)^2}{2} (1 - \cos 2\theta) + \frac{(R/b)^2}{2} (1 + \cos 2\theta) \right)^{-1/2} - 1 \tag{6}$$

By volume conservation, it can be shown that $R/a = (b/a)^{1/3}$ and $R/b = (b/a)^{-2/3}$.

$$\varepsilon = \left( \frac{(b/a)^{2/3}}{2} (1 - \cos 2\theta) + \frac{(b/a)^{-4/3}}{2} (1 + \cos 2\theta) \right)^{-1/2} - 1 \tag{7}$$

Applying double angle trigonometric identities and some algebra, we simplify to

$$\varepsilon = \left( \frac{b}{a} \right)^{2/3} \left( \sin^2 \theta \left( \frac{b}{a} \right)^2 + \cos^2 \theta \right)^{-1/2} - 1 \tag{8}$$

We can also express this as $\varepsilon = \alpha \left(\frac{b}{a}\right)^{2/3} - 1$, where $\alpha = \left(\sin^2 \theta \left(\frac{b}{a}\right)^2 + \cos^2 \theta\right)^{-1/2}$ serves as an angle correction factor. When $\theta = 0$, then $\alpha = 1$ and we recover the familiar limit $\varepsilon = \left(\frac{b}{a}\right)^{2/3} - 1$.

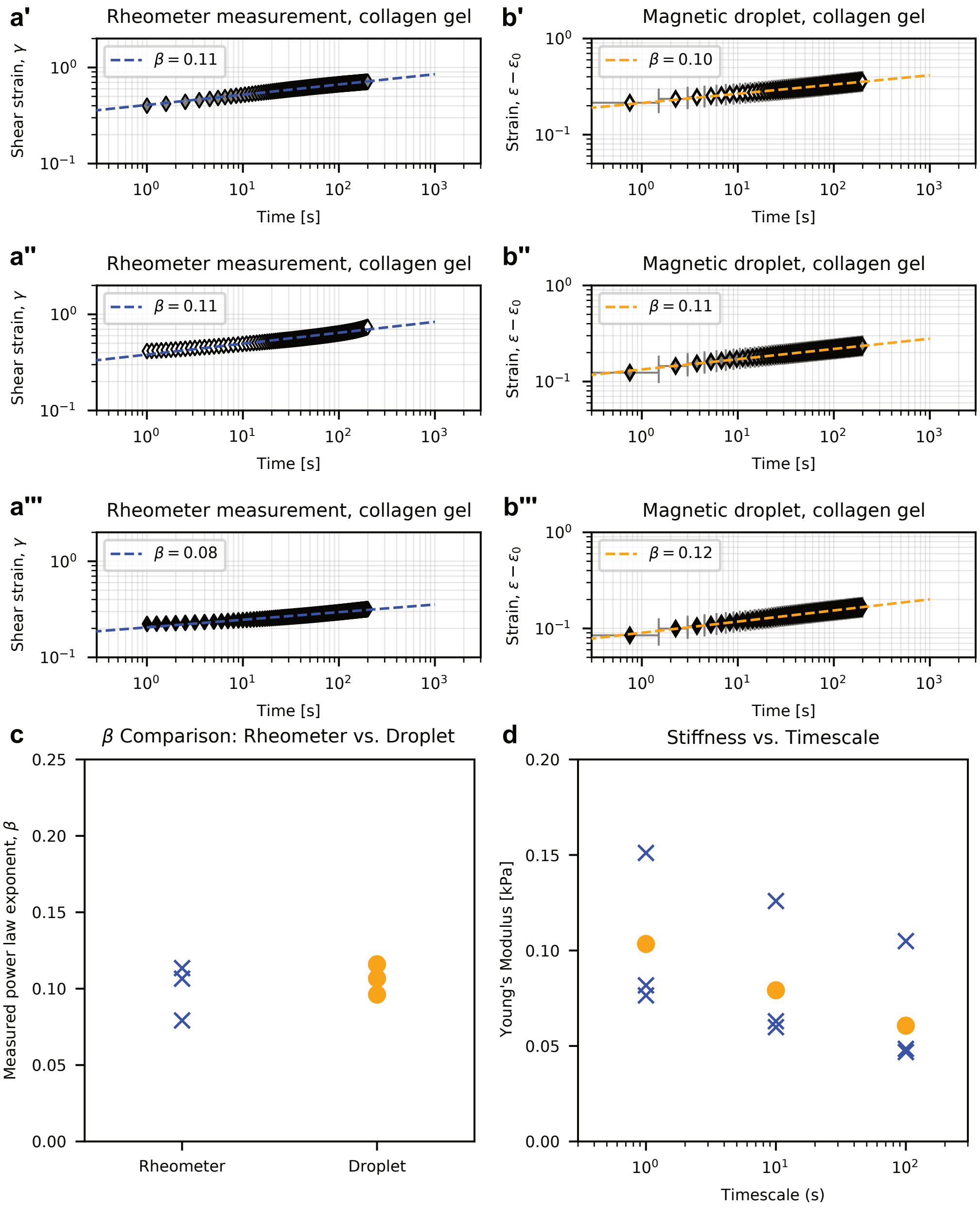

**Supplementary Fig. 6. Comparison of viscoelastic characterizations of collagen gels by magnetic droplet and shear rheometer show good agreement.** **a**, Shear strain versus time recordings for three creep tests of 1.76 mg/mL type I collagen gel on a shear plate rheometer, including power law fits shown as dashed lines. Scaling exponent $\beta$ is shown in legend. Shear stress $\tau = 10$ Pa. **b**, Strain $\epsilon$ minus pre-strain $\epsilon_0$ plotted against time for three creep tests of 1.76 mg/mL type I collagen gels with ferrofluid droplets deformed by magnetic stress. (b'-b") applied stress not calibrated; (b") applied stress $\sigma = 21.5$ Pa. **c**, Power law exponents obtained from rheometer (blue x) and droplet-based (orange circle) creep tests. **d**, Young's moduli of collagen gels computed at timescales of 1, 10, and 100 seconds from rheometer (blue x) and droplet-based (orange circle) data.

### 4 Supplementary Tables

#### 4.1 Supplementary Table 1

| $T$ [°C] | Visc. Std. | $N$ | $\eta_{exp}$ (mean ± s.e.m.) [Pa·s] | $\eta_{spec}$ [Pa·s] | Lot number |
|---|---|---|---|---|---|
| 25 | N62000 | 5 | $182 \pm 6$ | 185.800 | 16201 |
| | N190000 | 3 | $523 \pm 15$ | 523.300 | 14101d |
| 37 | N15000 | 4 | $17.8 \pm 0.7$ | | |
| | N62000 | 4 | $74.2 \pm 2.0$ | | |
| | N190000 | 3 | $215 \pm 7$ | | |
| | N450000 | 3 | $588 \pm 23$ | | |
| | N2700000 | 5 | $(2.05 \pm 0.07) \cdot 10^3$ | | |

**Supplementary Table 1: Measurements of viscosity standards on rheometer.** The viscosities of the viscous standards (Cannon Instrument Company) probed on a shear plate rheometer (Kinexus Ultra+, Malvern Instruments Limited). $T$: Temperature, Visc. Std.: Viscosity standard, $N$: Number of measurements, $\eta_{exp}$: mean experimental dynamic viscosity, s.e.m.: Standard error of the mean, $\eta_{spec}$: dynamic viscosity provided by the manufacturer for this lot number at temperature $T$ ($\eta_{spec}$ was only provided for $T = 37$°C). The standard error of the mean was calculated from the unbiased sample standard deviation.
